# Longitudinal MRI reveals hormone-dependent brain remodeling supporting preserved cognition in aged females

**DOI:** 10.64898/2026.04.02.715844

**Authors:** Egoa Ugarte-Pérez, Elena Espinós Soler, Antonio Cerdán Cerdá, Aroa S. Maroto, Patricia Martínez-Tazo, Maximilian Eggl, Laura Pérez-Cervera, Santiago Canals, Silvia De Santis

**Affiliations:** Instituto de Neurociencias CSIC-UMH, Sant Joan d’Alacant, Spain

## Abstract

Adaptive plasticity, the brain’s capacity to counteract structural decline and preserve performance with age, is a hallmark of successful aging, yet its biological underpinnings remain poorly understood. Here, we combined longitudinal magnetic resonance imaging (MRI) spanning the entire lifespan with electrophysiology, immunohistochemistry, and behavioral assays in female and male rats to identify key region- and systems-level mechanisms underlying this process. Resting-state functional MRI revealed a striking sex-specific pattern of connectivity reorganization in anterior brain regions, emerging in midlife and more pronounced in females. Microstructural MRI and histological analyses linked increased connectivity to prolonged white matter preservation and downstream maintenance of neuronal function in the female prefrontal cortex, while electrophysiological recordings demonstrated enhanced effective connectivity in the same region in aged females. Behaviorally, enhanced anterior connectivity was associated with superior memory performance. Ovariectomy at a critical time point for white matter maturation compromised this female-specific neuroprotection, disrupting microstructural integrity and functional reorganization, thereby highlighting the role of sex hormones in shaping these trajectories. Together, these findings identify a novel sexually dimorphic pattern of functional reorganization in anterior brain regions and point to estrogen availability during critical periods as a key modulator of brain aging.

## Introduction

As global life expectancy continues to rise, understanding how the brain adapts to structural decline to preserve function is essential for developing strategies that promote brain health across the lifespan. Increasing evidence suggests that healthy brain aging involves adaptive plasticity at the network level, whereby functional connectivity is reorganized to maintain cognitive performance despite structural decline^1–3^. While delayed microstructural deterioration is thought to support adaptive circuit reorganization and cognitive preservation during aging^4,5^, the cellular mechanisms and sex-specific determinants that govern these processes remain largely unknown. Aging affects neurobiological processes from the molecular and cellular levels to large-scale neural systems^6,7,8^. Advances in magnetic resonance imaging (MRI), one of the few techniques able to encompass these different scales, have greatly enhanced our understanding of brain changes across the lifespan. MRI studies show that aging is associated with reductions in grey matter volume^9^, while blood oxygen level–dependent imaging reveals functional alterations in regions critical for higher cognitive functions. In addition to connectivity reduction within several primary sensory and cognitive networks^10,11^, such alterations include age-related increases in frontal areas connectivity that correlate with better task performance in memory and decision-making tasks^12–14^, hence interpreted as a compensatory response. Diffusion-weighted MRI (dw-MRI) further demonstrates widespread microstructural alterations in white matter with advancing age^15^.

Recent work indicates that white matter changes may emerge earlier in males than in females^16^. Therefore, sex is increasingly seen as a key factor in the aging process, influencing brain structure, connectivity, and vulnerability to age-related diseases^17^. Multiple factors interact to shape these sex-specific patterns^18^. Among them, sex hormones, particularly estrogens, exert broad neuroprotective actions in the brain by regulating synaptic plasticity, cellular metabolism, and inflammatory pathways through estrogen receptor–dependent signaling cascades^19,20^. Addressing sexual dimorphism in aging is crucial; however, female underrepresentation in preclinical studies continues to limit our understanding^21,22^.

Rodent models, which recapitulate key features of human aging while compressing it into a tractable timescale, are essential for dissecting the cascade of events leading to cognitive decline and for probing the impact of modulatory factors such as sex hormones. However, despite confirming conserved age-related vulnerabilities in regions such as the frontal lobe and hippocampus^23,24^, most studies lack the longitudinal depth, lifespan coverage, or female representation needed to link aging trajectories to preserved cognition. Furthermore, translational gaps persist, as preclinical and clinical aging studies often employ disparate methodologies, complicating direct comparison and interpretation. In this study, we address these gaps using a systems biology approach. A dense longitudinal design tracking microstructural and functional trajectories across the lifespan reveals how specific brain regions adapt to age-related structural reorganization. We performed comprehensive longitudinal multimodal analyses in rats, including microstructural and resting-state functional MRI (rs-fMRI) across two years of the lifespan, alongside electrophysiology, immunohistochemistry, Monte Carlo simulations, and behavioral assessments at imaging-guided timepoints. Complementary analyses of microstructural MRI in human datasets strengthened the translational relevance of our findings. Using this approach, we uncover a novel sex-specific feature of aging: a midlife-emerging increase in functional connectivity localized to anterior brain regions, predicted by early-life white matter microstructure and markedly stronger in females. We further dissect the cellular, behavioral, and electrophysiological bases of this phenomenon, identifying sex hormones as key modulators of brain aging trajectories and shedding light on the strategies through which the brain adapts to aging.

## Results

Our first objective was to use longitudinal MRI to characterize brain structure, microstructure and function along the entire lifespan and in the whole rat brain. When examining longitudinal trajectories of both functional and microstructural MRI markers over two years of observation in rats (Fig. 1a), we found that our aging paradigm closely recapitulated key patterns observed in human imaging studies. Specifically, white matter microstructural indices such as fractional anisotropy (FA) and restricted water fraction (RF) - an index of integrity of the axon-myelin unit - exhibited distinct phases, with an initial period of maturation followed by a gradual decline (Fig. 1b-c). Model selection based on the Bayesian Information Criterion further revealed that most white matter regions were best fitted by segmented regression rather than linear models (11 out of 12 regions for FA and all 12 for RF, see Supplementary Table 1). Notably, when comparing across sexes, the breakpoint of these trajectories occurred significantly later in females, indicating a delayed onset of microstructural integrity decline (Fig. 1d). This delay was consistently observed for both FA and RF across all white matter regions (paired t-test: FA p=0.001, RF p=0.018). In white matter, the most plausible substrate underlying these patterns is myelin remodeling^25^. To test this, we quantified multiple histological markers in a subset of animals taken out of the longitudinal experiment aged 30–120 postnatal days (PND). Among all markers examined, myelin basic protein was the only one showing a significant age effect (Fig. S1). In addition, when comparing myelin trajectories across sexes for all white matter regions of interest, males exhibited a trend toward faster initial maturation (p = 0.06) and a significantly steeper decrease after 3 months (p = 0.032), consistent with earlier and more pronounced myelin remodeling (Fig. S1). More coarse metrics like brain size instead follows a linear trend with age, with no significant quadratic age or sex effect (Fig. S2).

**Figure 1.**
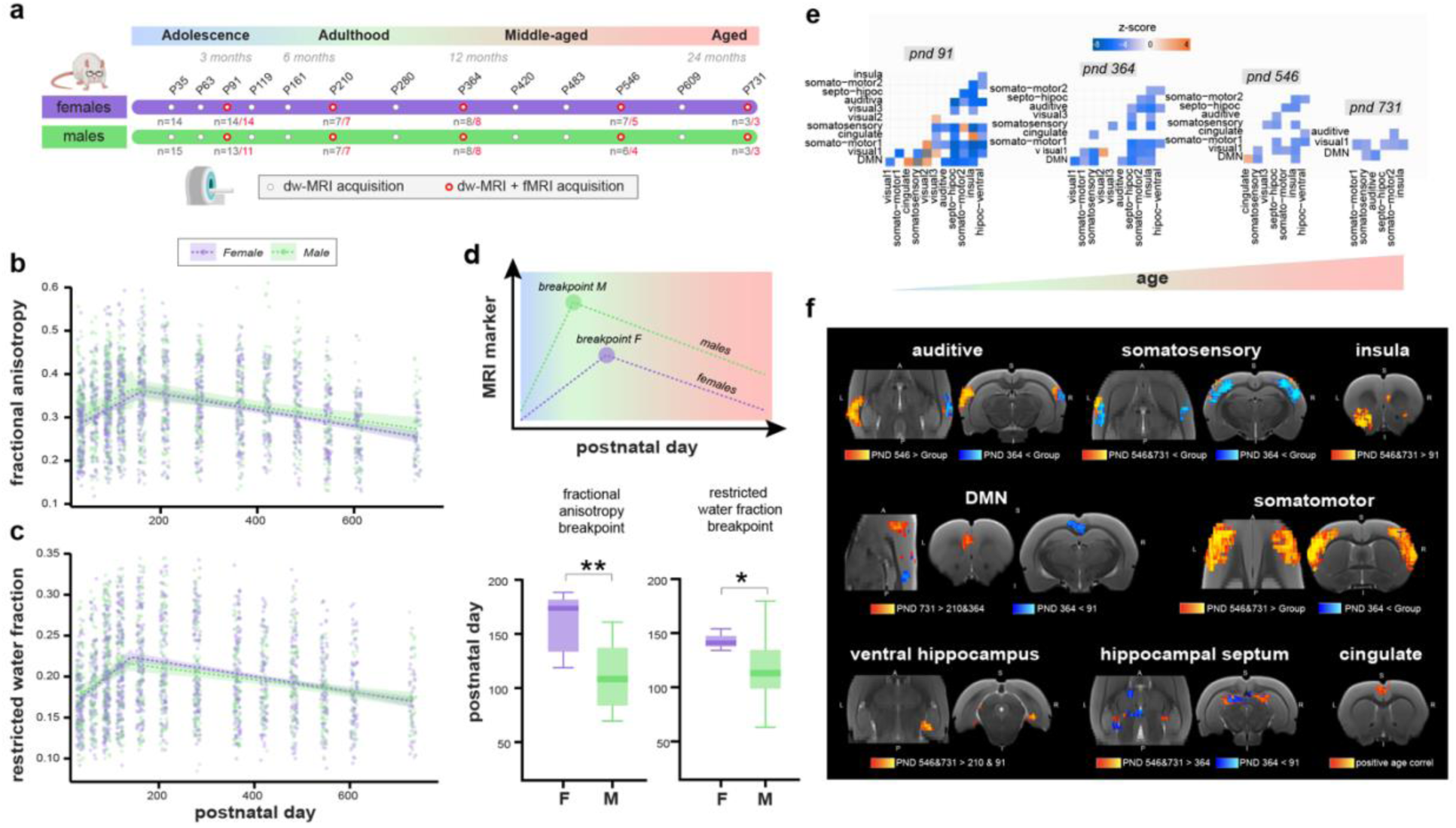
Longitudinal structural and functional MRI characterization across the lifespan. **a.** Experimental design of the longitudinal study. Dw-MRI was acquired at the ages indicated by white dots, whereas rs-fMRI acquisitions are marked by red dots. **b.** Longitudinal trajectory of FA across all white matter regions and animals. Solid lines indicate model fit for females (purple) and males (green), with shaded areas representing the 95% confidence bands. **c.** Same for RF. **d**. Upper panel: illustration of the segmented regression analysis to estimate sex-specific age-related breakpoints in MRI markers. The lower panel shows the average breakpoint across all white matter regions for FA (left) and RF (right); females are shown in purple and males in green, with breakpoints occurring significantly later in females for both markers (paired t-test: FA p = 0.0011, RF p = 0.0179). Bar plots represent median ± standard errors. **e.** L2-regularized partial correlation between resting-state networks for different ages, reported as z-scores quantifying the strength of correlations between pairs of significantly correlated networks. **f**. Comparison of within-network functional connectivity across ages. Blue indicates clusters showing decreasing connectivity with age, whereas red indicates clusters showing increasing connectivity. Abbreviations: DMN = default mode network; Septo-hipoc = septo-hippocampal network; F = female; M = male.

In functional timeseries during resting state, independent component analysis combined with dual regression allowed us to assess both between-network and within-network connectivity. Between-network connectivity was defined as the strength of the correlation between network-specific timecourses, whereas within-network connectivity was assessed voxel-wise as the contribution of each voxel to the average network timecourse. When between-network connectivity was examined, an age-related decrease in the number of significant connections was observed, with a preferential loss of positively connected nodes (Fig. 1e, ICA-derived networks are shown in Fig. S3). However, within-network connectivity showed a more complex pattern, with both increases and decreases depending on age and the specific subnetwork, as shown in Fig. 1f. Several networks exhibited bilateral reductions in connectivity until one year of age, followed by bilateral increases between one and two years. No significant effects of age or sex were observed in the weighted average BOLD signal of any network.

Next, we examined the increase in within-network connectivity in greater detail. Voxel-wise mapping of age-related effects within the default mode network (DMN), a conserved network spanning anterior and posterior regions in both rodents and humans, revealed that this increase was preferentially localized to anterior regions (Fig. 2a). For this network, a significant interaction between sex and age also existed (Fig. 2a, light violet), localized in the anterior part. Examination of average connectivity estimates across timepoints demonstrated that old females showed significantly higher within-network connectivity in anterior regions compared to young females (unpaired t-test, p = 0.007), while no significant effect was observed in males (Fig. 2b). Notably, when testing for a sex × age interaction on the observed increase in within-network resting-state functional connectivity across all networks, significant voxel-wise clusters emerged exclusively in anterior areas of the somatomotor, DMN, visual, cingulate, and somatosensory networks (Fig. 2c).

**Figure 2.**
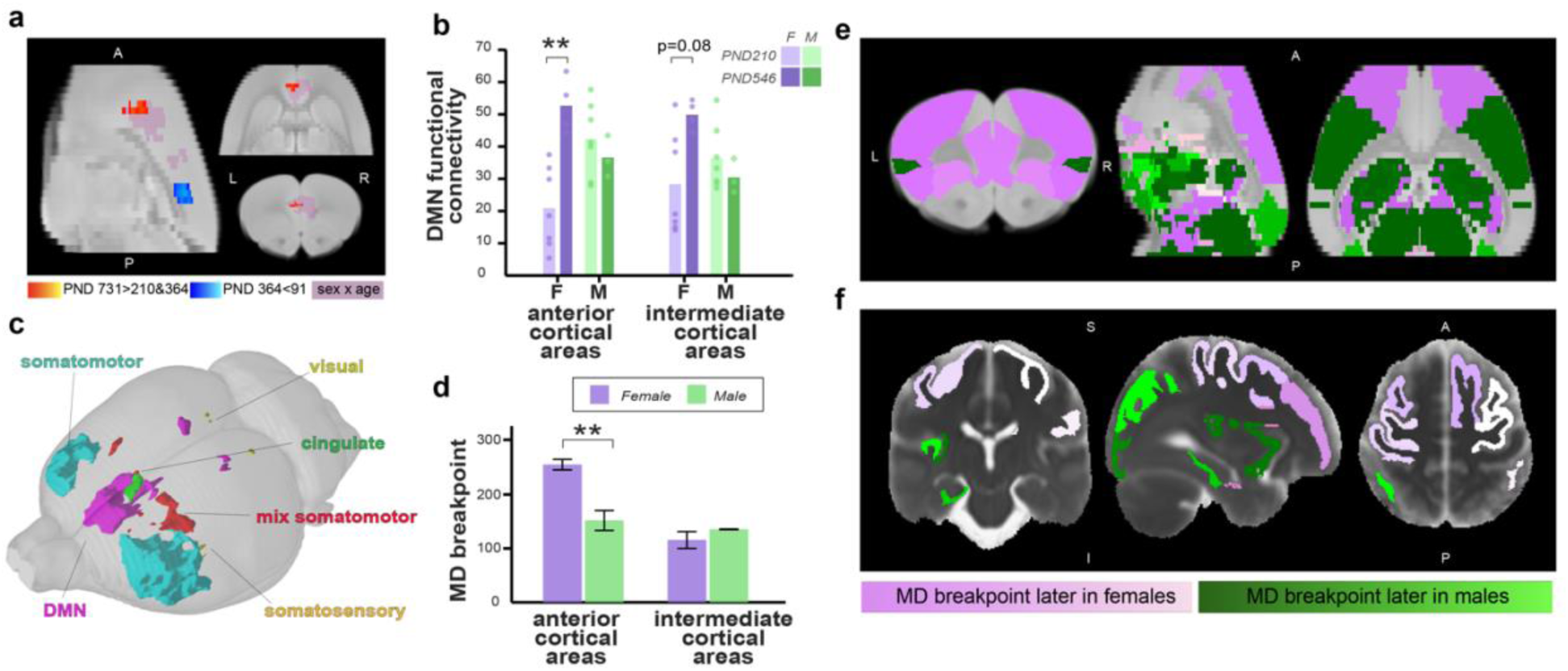
Sexually dimorphic anterior connectivity reorganization and delayed microstructural integrity decline in female anterior cortical areas. **a**. In the DMN, clusters showing a significant decrease in functional connectivity in the age range 0-400 PND are shown in blue, whereas clusters with significant increase in the 500-700 PND range are shown in red. In the same network, significant sex x age interactions are shown in pink shadow. **b.** Within-network functional connectivity in males (green) and females (purple), averaged across voxels belonging to anterior and posterior DMN, and for different ages (light color, PND 210, dark color, PNS 546). In females, connectivity is significantly greater in anterior cortical areas at PND 546 (dark purple) compared to PND 210 (unpaired t-test, p = 0.007), with a trend for increased connectivity in intermediate cortical areas (unpaired t-test, p = 0.08). No significant differences are observed in males. **c**. Of all RSNs (somatomotor, visual, cingulate, somatosensory, and DMN), the clusters displaying a significant sex × age interaction are localized exclusively in anterior areas. **d**. Averaged MD breakpoint in males (green) and females (purple) for anterior versus intermediate cortical areas, showing a delayed inflection point in anterior cortical areas only (unpaired t-test, p = 0.005). Error bars represent standard errors. **e**. Difference in the age at which the MD breakpoint is observed between sexes (purple, breakpoint earlier in males; green, breakpoint earlier in females) shown in regions with significant quadratic trajectories with age. **f**. Same as e, but for the human cohort of ref. 16.

Turning to microstructural MRI in grey matter, we observed that mean diffusivity (MD) versus age showed more heterogeneous trajectories when compared to the white matter trajectories (Supplementary Table 2). However, when comparing MD breakpoint values between anterior cortical areas (infralimbic, prelimbic, motor, and orbital cortices) and intermediate cortical areas (somatosensory and parietal association cortices), the MD breakpoint was significantly delayed in females only in anterior regions (unpaired t-test, p = 0.005, Fig. 2d). This was further illustrated by mapping the difference in days between female and male breakpoints (Fig. 2e).

Finally, we re-analyzed the cross-sectional human MRI dataset from ref. ^16^, which originally focused on white matter biomarkers, to examine MD in grey matter. Consistent with our rat data, regions exhibiting a delayed MD breakpoint in females were predominantly located in anterior cortical areas (Fig. 2f).

By leveraging the longitudinal design, we subsequently delineated the temporal cascade of age-related structural, microstructural and functional changes. Temporally, the sex effect first emerged at the structural level within white matter, with females showing preserved microstructural integrity due to a significant delayed inflection point in FA (Fig.1d) and in the average slope of the MD post-breakpoint trajectories (paired t-test, p=0.04), starting around 100 PND (Fig. 3a). Although the sex effect was not restricted to anterior white matter breakpoint, the post-breakpoint slope in MD was significantly lower in females specifically within this region, as evidenced by the correlation between the slope and the anteroposterior coordinate (Pearson correlation; r= 0.63, p=0.029) (Fig. 3b), and which was not significant in males (Fig. S4a). Subsequently, MD reaches the breakpoint in grey matter, with a sex effect localized to anterior cortical areas, indicating early breakpoint in males (100 post-natal days vs 250, Fig. 3c). Increases in within-network functional connectivity emerged concurrently with grey matter remodeling in females, beginning around 250 PND, whereas in males they occurred substantially later (around 400 PND) (Fig. 3d). A schematic summary of the temporal sequence of these sex-specific effects is provided in Fig. 3e.

**Figure 3.**
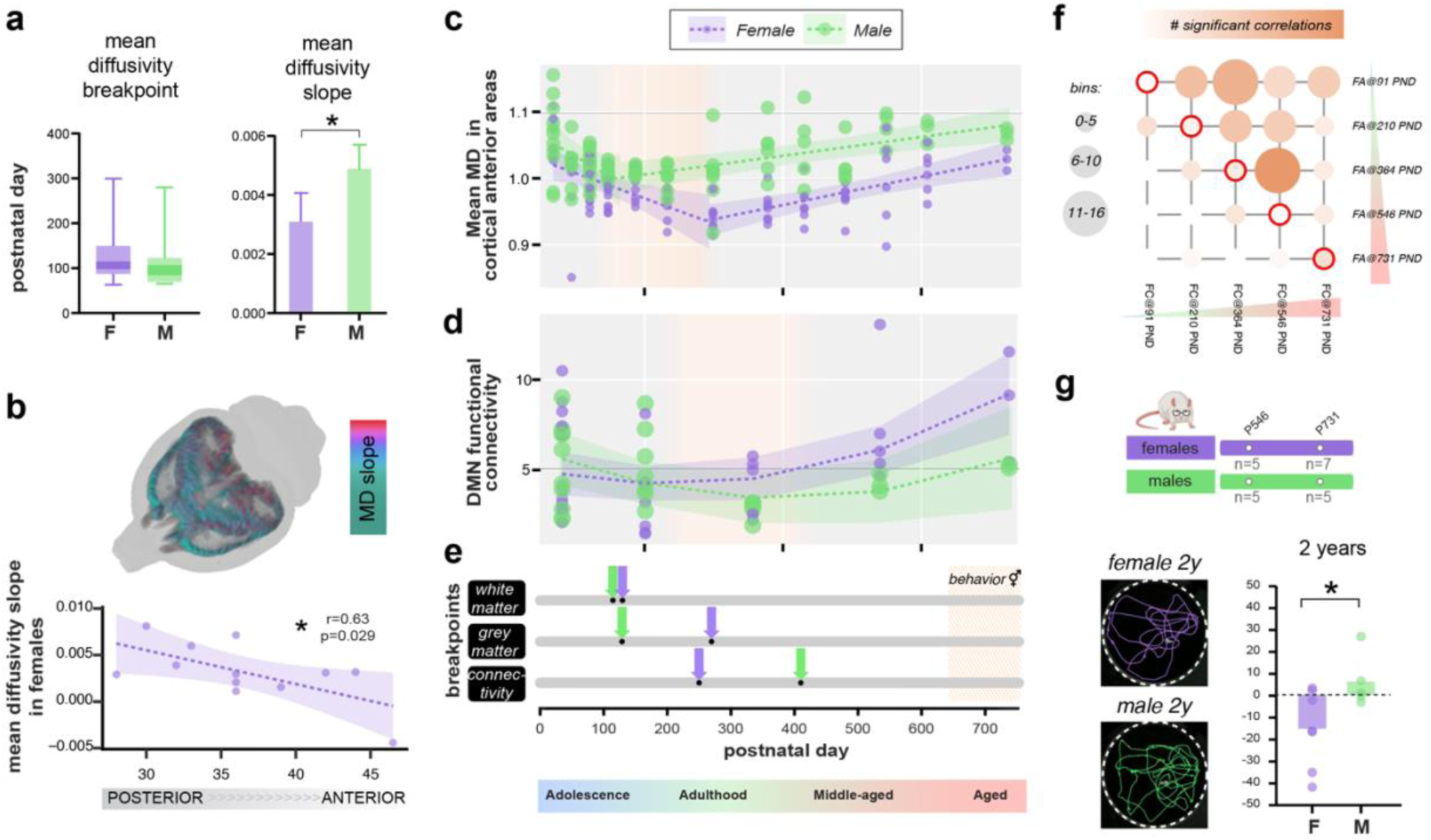
Temporal sequence and cognitive correlates of sex-specific microstructural integrity and functional connectivity reorganization across the lifespan. **a**. Mean breakpoint (left) and slope after breakpoint (right) of MD in white matter for female (purple) and male (green) rats (paired t-test, p=0.04 for slope). **b.** MD slope after breakpoint in white matter is plotted as a function of the anterior-posterior coordinate for females (Pearson correlation test, r= 0.63, p=0.029). The shaded area indicates the 95% confidence band. **c.** Mean MD trajectory in anterior cortical areas (infralimbic, prelimbic, motor, and orbital cortices) versus age. The breakpoint in female (purple) is around ∼270 PND, while in males around ∼130 PND (green), as assessed using a segmented regression model (discontinuous line). Shaded areas represent the 95% confidence bands. **d**. Functional connectivity in the DMN versus age and fitted quadratic model, showing a breakpoint around 250 days in females (purple) and around 410 days in males (green). Shaded areas represent the 95% confidence bands. **e**. Schematic summary of the temporal sequence of sex-specific microstructural and functional changes across the lifespan, highlighting the progression from white matter integrity to grey matter alterations and functional connectivity remodeling (females in purple, males in green). The orange square indicates the age range where significant differences in long-term memory are observed between sexes. **f**. Significant cross-modal correlations between average FA values in white matter and functional connectivity measures for all age pairs (Wilcoxon signed-rank test, p = 0.002). **g**. Experimental design of behavioral measures, water maze trajectories for one female and one male rats, and group differences for long-term memory at 2 years (unpaired t test, p = p=0.032). Abbreviations: PND = post-natal day; F = female; M = male; 2y = 2 years.

Cross-modal analyses across timepoints confirmed that early microstructural alterations predicted subsequent fMRI changes, indicating that preserved structural integrity preceded age-related remodeling in functional connectivity during aging. Specifically, correlations between mean white matter FA and within-networks connectivity across different networks revealed that higher FA at earlier timepoints was significantly associated with higher connectivity at later timepoints, as evidenced by a greater number of significant correlations above the diagonal (Wilcoxon signed-rank test, p = 0.002, Fig. 3f). A similar, albeit weaker, temporal relationship was observed between MD in grey matter regions and within-networks connectivity, but the contrast did not reach significance (Fig S4b).

We then tested whether the increased functional connectivity observed in anterior areas in females in the longitudinal cohort was associated with improved cognitive performance, thereby allowing us to interpret it as an effective compensatory mechanism. Indeed, in animals aged 2 years, females exhibited a significant faster long-term learning rate in the Morris Water Maze task compared to males, as indicated by the slope of latency to find the platform across five training days (unpaired t-test p=0.032) (Fig. 3g). No significant sex differences were observed at 1.5 years of age, nor in short-term memory or in motor function (Fig. S3 c and d, respectively).

Next, we examined microstructural changes in regions showing increased connectivity specifically in females. In white matter, decreases in FA and increases in MD are commonly interpreted as markers of declining microstructural integrity associated to myelin reorganization, whereas their biological substrate in grey matter remains less well established. Hence, we turned to age-related trajectories of advanced MRI markers in grey matter, such as RF and fiber dispersion (FD). In grey matter, these parameters are interpreted as neurite density and dispersion, respectively, and they can refer to both neurons and glia. While neurite density consistently declined with age across regions, neurite dispersion was selectively maintained or increased in anterior areas, such as the prelimbic or motor cortices (Fig. 4a). To further interpret these changes, we performed Monte Carlo simulations modeling the effects of neuronal, microglial, and astrocytic alterations on these MRI parameters. The pattern observed in anterior ROIs, i.e., decreased neurite density and increased dispersion, was most consistent with predominant neuronal alterations (1 sample Wilcoxon signed rank test, Benjamin-Hochberg corrected; p=2.7e-5 for neurite dispersion in neurons; p=1.9e-6 for neurite density for all three conditions after correction; Fig. 4b).

**Figure 4.**
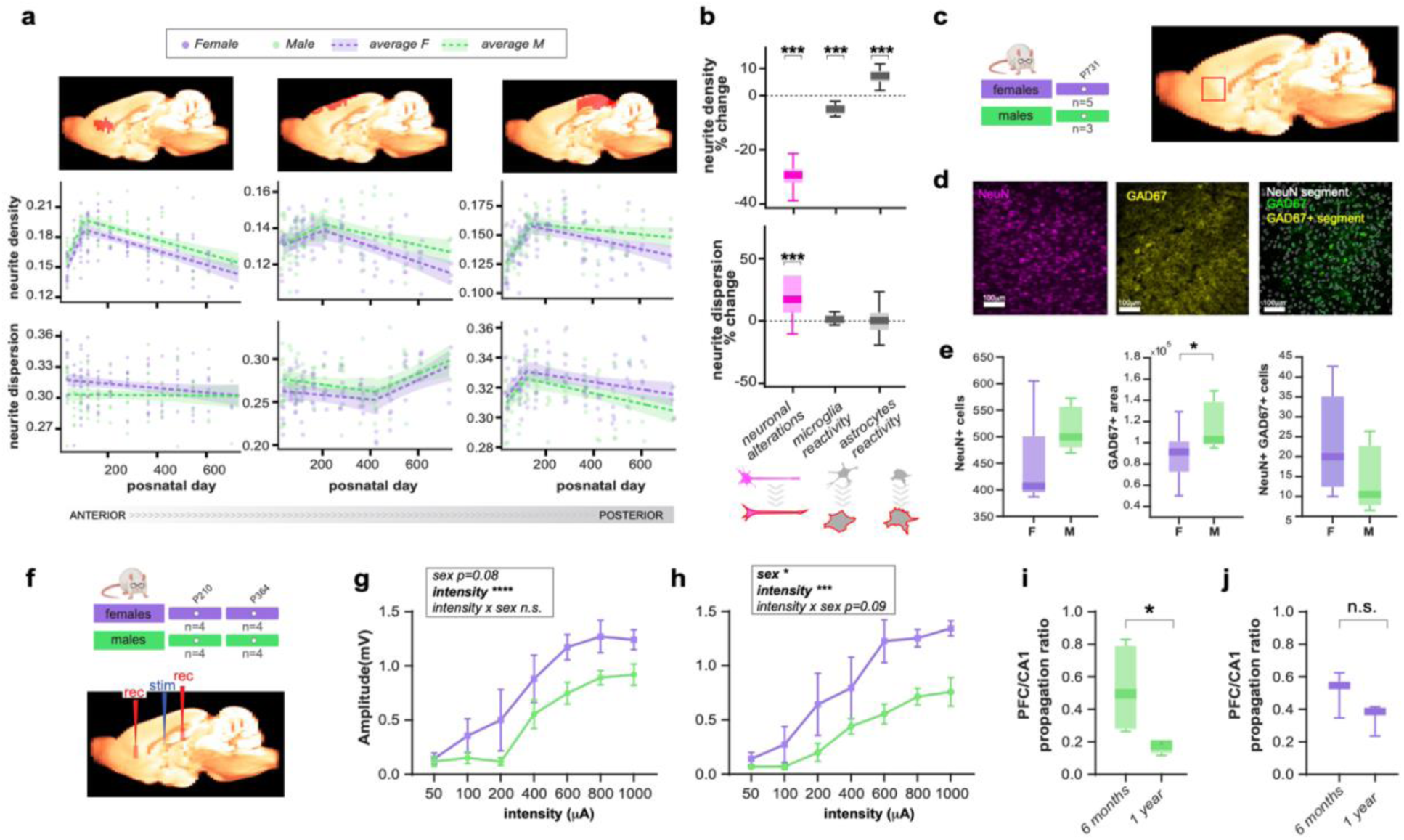
Cellular and electrophysiological correlates of functional connectivity remodeling in anterior regions from advanced dw-MRI, immunohistochemistry, and local field potential recordings. **a.** Age-related trajectories of neurite density and dispersion in anterior and posterior representative ROIs: prelimbic cortex at the left, motor cortex in the middle and retrosplenial cortex at the right. Segmented lines represent the best model fits (female in purple, male in green), and shaded areas indicate 95% confidence bands. **b.** Monte Carlo simulations of different substrates associated with neuronal, microglia or astrocyte alterations. For each altered cellular substrate, the upper graph shows predicted changes in neurite density, and the lower graph shows predicted changes in neurite dispersion. Data are shown as mean ± standard error**. c.** Immunohistochemistry experimental design (left), and the prelimbic cortex selected for analysis (right), highlighted with a red square. **d.** NeuN+ cells, GAD67 staining, and segmentation of GAD67+ neuronal cells in two representative animals. **e.** NeuN cell count (left, unpaired t-test, p=0.23), GAD67+ (center, unpaired t-test, p=0.04) and GAD67+ NeuN cell count (left, unpaired t-test, p=0.29) for regions-of-interest in prelimbic cortex, compared across sexes**. f.** Experimental scheme for the electrophysiological experiment. Top figure: ages and number of animals from a separate cohort used for the input/output experiment (female in purple, male in green). Bottom figure: schematic of recording sites in the prefrontal cortex and hippocampus, and stimulation in the fimbria. **g.** Evoked activity versus stimulation intensity in the prefrontal cortex at 6 months of age. (Repeated measures two-way ANOVA, main effect of intensity p < 0.0001; sex main effect, p = 0.08; female in purple, male in green). Data are shown as mean ± standard error. **h.** Same as g at 1 year of age. Repeated measures two-way ANOVA, intensity main effect, p < 0.0001; sex effect, p = 0.03; intensity × sex interaction, p = 0.09). Data are shown as mean ± standard error. **i-j**. Propagation ratio at different ages, shown separately for males (green; unpaired t-test across ages, p=0.04) and females (purple; p=0.18). Data are shown as mean ± standard error. Abbreviations: rec = recordings; stim = stimulus.

Immunohistochemical analyses at the final time point revealed no significant sex differences in neuronal or non-neuronal counts (Fig. 4e left for prefrontal cortex, and Fig. S5 for other markers and a more lateral histological slice). In contrast, quantification of inhibitory markers in prefrontal cortex showed that GAD67 staining, which quantifies GABAergic signaling, was significantly increased in males, indicating a higher density of inhibitory terminals (unpaired t-test, p = 0.04; Fig. 4e). However, no sex differences were observed in GAD-positive neuronal cell counts (Fig. 4e, left).

Next, we sought to test the electrophysiological basis of the increased connectivity observed in aged animals. In a new cohort, we performed local field potential recordings in the prefrontal cortex (prelimbic/infralimbic) and the hippocampus while stimulating the fimbria, which provides the majority of axonal projections from the hippocampus to the prefrontal cortex. Stimulus-response analysis demonstrated significant enhanced evoked responses in the female prefrontal cortex, measured as the amplitude of the evoked potential across different stimulation intensities, starting from 1 year of age. At 1 year of age, repeated measures two-way ANOVA revealed significant main effects of sex (p = 0.03), with a trend toward a sex × intensity interaction (p = 0.09), indicating increased effective connectivity within this pathway. At 6 months the same analysis did not reach significant sex differences, showing only a trend toward a sex effect (p = 0.08). The evoked response in the hippocampus was also analyzed using the same statistical approach. At 6 months of age, repeated measures two-way ANOVA revealed no significant effect of sex or sex × intensity interaction (Fig. S6). At 1 year of age, no main effect of sex was detected, but a significant sex × intensity interaction emerged (p = 0.049, Fig. S6), however, no significant effects in post-hoc comparisons were found. Importantly, normalizing the polysynaptic evoked response in the prefrontal cortex by the activation magnitude in the hippocampal CA1 region, thus computing an effective propagation ratio CA1→prefrontal cortex, revealed an age-related decrease in the efficiency of this connection in males (unpaired t-test, p=0.03, Fig. 4i), but not in females (Fig. 4j), further supporting preserved communication in aged female rats.

Finally, to test the influence of sex hormones on aging trajectories and the emergence of increased functional connectivity with age, we analyzed a new cohort of animals in which ovariectomy (OVX) was performed at 2 months of age, during the critical window for white matter integrity identified by our longitudinal analyses, and compared them with age-matched sham-operated controls (scheme in Fig. 5a).

**Figure 5.**
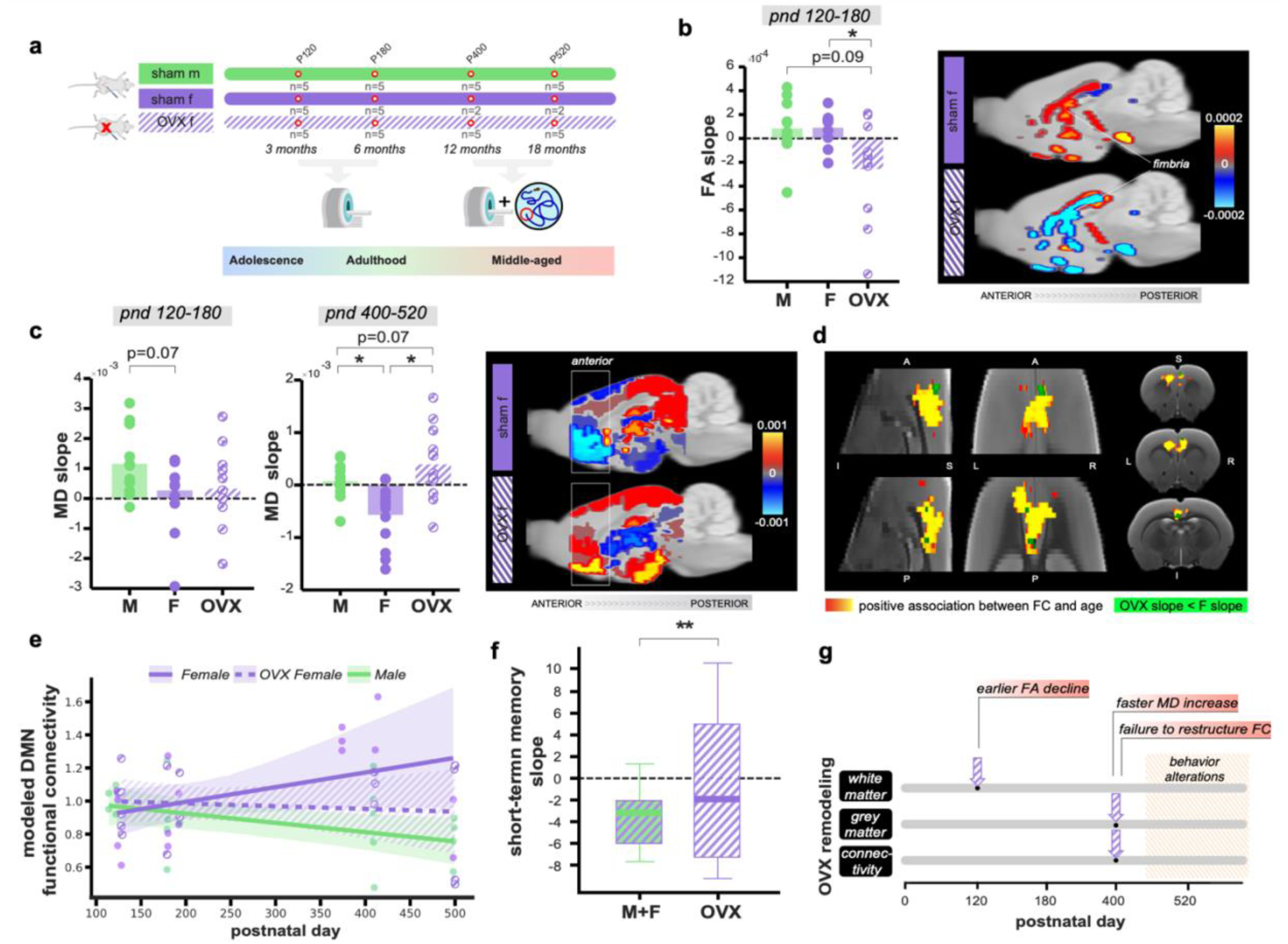
Premature loss of sex hormones accelerates aging trajectories in females. **a**. Experimental design comparing OVX animals with sham-operated female and male rats. Sham males are shown in green, sham females in purple, and OVX animals in purple and white stripes. **b.** FA slope between 3 and 6 months for all white matter regions. Repeated-measures ANOVA (F(2,20)=6.19, p=0.008) followed by post-hoc comparisons (OVX vs. F p=0.02 FDR-corrected, OVX vs. M p=0.09 FDR-corrected) indicate lower slope in OVX animals (left panel). In the right panel, the slope is projected onto the white matter parcellation for female sham (above) and OVX animals (below). **c**. MD slope between 3 and 6 months (left) and between 1 and 1,5 years (middle) compared across groups. Repeated-measures ANOVAs show no significant group effect between 3 and 6 months (F(2,24)=3.37, p=0.051) but significant group effect between 1 and 1,5 years F(2,24)=9.32, p=0.001). OVX animals have steeper MD slopes than female sham animals (p=0.016 corrected) and a trend towards steeper MD slopes than male sham animals (p=0.07 corrected). In the right panel, the slope between 1 and 1,5 years is projected onto the grey matter parcellation for female sham (above) and OVX animals (below). **d**. DMN voxels with significant positive age effect in within-network functional connectivity are shown in red, while in green, voxels where the slope with age is lower in OVX animals compared to sham females are shown in green. **e**. Within-network functional connectivity, reported as z-scores across ages and groups, showed with confidence intervals corresponding to the standard deviation. **f**. Short-term memory slope in water maze task at 1.5 years for OVX vs sham animals (unpaired t-test, p=0.002). Sham animals were grouped together, as no differences were observed at this age, consistent with the longitudinal experiment. **g.** Summary of the temporal sequence of sex-specific microstructural and functional changes across the lifespan in OVX animals. Abbreviations: Sham m = sham male; sham f = sham female; OVX f= ovariectomized female; F = female; M = male.

When comparing FA slopes across white matter regions in animals between 3 and 6 months of age, we found a significant group effect (repeated-measures ANOVA, p=0.008; Fig. 5b). Post-hoc comparisons revealed that OVX animals reached the inflection point of white matter trajectories significantly earlier than sham females (OVX vs female p=0.02 corrected) and showed a similar trend compared with males (OVX vs male p=0.09 corrected).

Analysis of MD slopes across anterior grey matter regions between 3 and 6 months showed a trend toward a group effect (repeated-measures ANOVA, p=0.051; Fig. 5c left). However, between 12 and 18 months a strong group effect emerged (repeated-measures ANOVA, p=0.001; Fig. 5c right). Post-hoc comparisons confirmed that OVX animals exhibited steeper MD increases than sham females (p=0.016, FDR-corrected) and a trend toward steeper slopes compared with males (p=0.07, FDR-corrected). Similar analyses in intermediate grey matter regions revealed a significant group effect (repeated-measures ANOVA, p=0.02), although post-hoc comparisons were not significant. Together, these results indicate an earlier onset of grey matter microstructural decline in OVX animals, primarily affecting anterior brain regions.

We next examined functional connectivity within the anterior DMN. In sham animals, we replicated in this cohort the increased functional connectivity with increasing age (yellow cluster of p<0.05 voxels, Fig. 5d). Importantly, when comparing the slope versus age, several voxels show significant lower slope in OVX animals compared with sham females (green cluster of p<0.05 voxels, Fig. 5d). This is directly plotted as a z-score in Fig. 5e. Finally, behavioral analysis revealed significant differences between OVX and sham animals in the slope of short-term memory performance in the water maze at 1.5 years (unpaired t-test, p=0.002; Fig. 5f), indicating impaired cognition in OVX animals. This contrast was not significant at 1 year (p=0.32), suggesting that behavioral deficits emerge after the loss of white matter integrity and the failure of functional connectivity remodeling, as summarized in Fig. 5g. No significant differences in long-term memory were observed between sham and OVX animals.

## Discussion

In this study, we dissect the cellular, electrophysiological, and behavioral correlates of functional remodeling in a rat model of aging, revealing its spatial localization, temporal sequence, and distinct mechanistic signatures across sexes, including ovariectomized animals. This was achieved in three steps. First, longitudinal MRI across the lifespan enabled an unbiased identification of sex-specific critical periods and vulnerable brain regions. Second, we performed detailed electrophysiological and morphological analyses at key imaging-derived time points. Finally, we examined the contribution of sex hormones to the observed patterns.

### Age-driven brain plasticity in human and rodent aging

Adaptive network plasticity in aging has been studied largely in humans^26^, but several limitations remain. First, few studies span a sufficiently wide age range to capture the point at which compensatory processes may emerge^27^. Second, most aging studies are cross-sectional, preventing direct measurement of individual trajectories. Finally, it remains debated whether increased connectivity truly represents a compensatory mechanism. Evidence remains mixed, with some studies linking increased connectivity to improved performance^13^, whereas others associate it with less efficient processing or declining function^28^.

While these challenges are, in principle, addressable in animal models, establishing convincing models of aging has remained difficult. Most rodent fMRI studies rely on seed-based approaches that focus on predefined regions rather than whole-brain organization^29,30^, or span only a portion of the lifespan^31,32^. Here, through dense multimodal MRI data collected over two years, we show that aging is accompanied by the emergence of functional connectivity remodeling associated with preserved cognitive performance. Importantly, our longitudinal analyses reveal a striking convergence between rodent and human aging across multiple MRI modalities. In both species, white matter microstructure follows a biphasic trajectory, with an early phase of maturation succeeded by a gradual decrease, a pattern consistently reported in human lifespan neuroimaging studies^33,34,35,36,37^ and primarily attributed to myelin maturation and remodeling^38^.

We show that sex significantly shapes MRI trajectories, matching observations in large human cohorts showing that sex mediates aging effects on white matter microstructure integrity^39,40^. Importantly, we replicate in rats our previous finding that the inflection point of age-related structural trajectories occurs later in females than in males^16^, reinforcing the translational relevance of the model and then use of metrics like the inflection point, which require high temporal sampling across the lifespan.

At the functional level, we observed reductions in between-network connectivity with advancing age. fMRI findings in humans suggest that aging is associated with a complex pattern of neural activity changes, involving both increased and decreased activation in older compared with younger individuals^41,42,43^. Nonetheless, there is a relative consensus that certain regions present an age-related increase of brain activity^44^. Indeed, in our data within-network analyses revealed localized increase in connectivity emerging at later ages, particularly in anterior regions of the default mode and prefrontal networks. Importantly, we were able to replicate this finding in a second experiment with a different cohort. This pattern aligns well with previous fMRI evidence of compensatory recruitment in frontal regions during cognitive aging^45^ and studies indicating that long-range connectivity decreased with age whereas short-range connections get stronger^46^. However, our study critically links the connectivity remodeling to the emergence of a sexual dimorphism. The prefrontal localization of the sex effect aligns with known sex differences in aging of frontal areas in more coarse measures. Large neuroimaging cohorts (e.g., ENIGMA Lifespan, UK Biobank) show slower frontal cortical thinning and smaller age-related volume loss in women, particularly in dorsolateral prefrontal cortex^39^. FMRI studies also indicate that older women maintain stronger prefrontal cortex activation during executive and working memory tasks^47^. Importantly, in our study we support and extend these fMRI observations using electrophysiological recordings, demonstrating sex-dependent neuronal activity in the prefrontal cortex, particularly evident from one year of age, together with histological evidence indicating differences in inhibitory markers across sexes.

### Structural alterations drive functional changes in aging

Examination of the temporal sequence across our multimodal analyses reveals that early microstructural alterations in white matter precede grey matter remodeling and the increased within-network connectivity, that ultimately culminates in cognitive preservation. This interpretation is further supported by cross-modal correlations showing that fractional anisotropy at earlier timepoints, an index of myelination, predicts functional connectivity at later stages. Indeed, several studies underscore the importance of structural maintenance in aging^48^, and suggest that increased activity in the anterior cortical areas may be triggered by age-related frontal grey matter atrophy^49^. Further supporting the link between structure and function is the observation of higher evoked responses in the female prefrontal cortex. In addition, reduced GAD expression in females suggests a shift in the local excitation/inhibition balance towards excitation, which likely contributes to the enhanced evoked responses. Together, our histological and electrophysiological findings suggest that the observed within-network functional remodeling arises from two coexisting mechanisms: (i) delayed deterioration of white matter integrity preserving effectiveness of transmission, and (ii) higher excitation/inhibition balance in females. Based on the temporal sequence revealed by our longitudinal analyses, we further propose that the former may at least partially drive the latter, setting the stage for progressive neuronal adaptations. Prolonged preservation of myelin in females, evidenced by imaging microstructural metrics and myelin immunostaining, may help sustain efficient and temporally coordinated axonal conduction, thereby contributing to circuit stability and within-network functional connectivity over time^50,51^. In contrast, earlier myelin deterioration may compromise axonal integrity and promote cortical “disconnection” and decreased excitatory capability in regions such as the prefrontal cortex^52,53^. Beyond its structural role, myelin integrity may also contribute to axonal maintenance through oligodendrocyte-mediated metabolic support, suggesting that age-related declines in myelin preservation and remyelination capacity could favor progressive axonal and neuronal vulnerability^51,54,55^. Consistent with this framework, the higher excitation/inhibition balance observed in aged females may be a consequence of prolonged structural integrity, and instrumental for successful network adaptation.

### Sexually dimorphic functional remodeling in aging

While the involvement of the anterior structures in aging is well established, less is known about how sex shapes these age-related patterns. Our most striking finding is a robust interaction between sex and aging that is spatially confined to anterior cortical regions. Notably, areas exhibiting higher within network connectivity in aged females overlap with regions showing delayed microstructural integrity decrease, particularly within the anterior cortical areas. Consistent with this, electrophysiological recordings reveal enhanced evoked responses in prefrontal cortex in females beginning at one year of age. This enhancement appears to be locally generated, as the magnitude of input to the region remains unchanged. We interpret these converging functional and structural changes as a compensatory mechanism rather than a nonspecific alteration in connectivity, as aged females outperform males in behavioral tasks that critically depend on anterior cortical areas functions.

Importantly, we uncovered a critical temporal relationship between compensatory processes and the onset of microstructural integrity changes. In females, the reorganization of connectivity is tightly synchronized with the onset of grey matter microstructural integrity changes. In contrast, males exhibit an earlier start of microstructural integrity changes in both white and grey matter, with increased connectivity arising at a delayed and desynchronized time point, ultimately leading to poorer performance on tasks mediated by anterior cortical areas. Together, these findings suggest that the connectivity increase in females may represent an adaptive response to maintain network efficiency as structural integrity begins to deteriorate.

### Mechanisms of anterior cortical areas remodeling in female and role of estrogens

Notably, the preferential localization of the sex effect to anterior cortical areas suggests a contribution of sex-hormone–dependent neuroprotective mechanisms. Our findings suggest that females have prolonged preservation of white matter microstructure, consistent with estrogen-supported myelin maintenance^56^, which delays the onset of grey-matter degeneration and preserves neuron function. The prefrontal cortex specificity may be related to larger amount of estrogen receptors in frontal white matter^57^. Another complementary mechanism may involve reduced dopamine clearance via catechol-O-methyltransferase (COMT) ^58^. Estrogens inhibit COMT, elevating extracellular dopamine and enhancing neuronal excitability^59^, particularly in the prefrontal cortex where COMT is the dominant clearance pathway^60,61^. This mechanism is consistent with our electrophysiological findings of increased neuronal responses in females, and with our immunohistochemistry findings of lower GABAergic terminals.

Given these results, we directly tested the contribution of estrogen availability to sex-specific aging trajectories using females ovariectomized during the critical window for white matter integrity identified by our longitudinal analyses. Ovariectomy accelerated white matter degeneration, followed by earlier and faster gray matter changes and a delayed emergence of functional reorganization, ultimately leading to poorer performance in water maze tasks. Previous studies have shown that ovariectomy and estrogen deprivation impair spatial and hippocampal-dependent memory in rodents and exacerbate cognitive decline^62,63^, whereas estrogen replacement can partially restore these deficits^64^. However, a key novelty of our study lies in linking sex hormone availability to longitudinal microstructural trajectories and adaptive network plasticity. These findings emphasize the relevance of estrogen availability during critical periods of brain maturation for shaping brain aging trajectories and provide a mechanistic basis for observational human studies reporting similar associations^65,66^.

### Limitations

This study has several limitations. In particular, the sample size at later time points was reduced due to animal loss. While this is an inherent challenge in longitudinal studies, it may have contributed to the reduced detection of significant between-network correlations. A general limitation of fMRI studies in aging is that BOLD signal changes can be influenced by age-related alterations in neurovascular coupling, potentially confounding the interpretation of functional connectivity differences. However, analyses of BOLD signal properties (in the present study as temporal variability) did not reveal significant age-related differences, supporting the interpretation that the observed functional changes are not solely driven by vascular effects. Importantly, this limitation is further mitigated by our multimodal approach, which combines fMRI with electrophysiological recordings that provide a direct measure of neuronal activity.

## Conclusions

In conclusion, our findings identify functional adaptive plasticity as a sex-specific process rooted in preserved brain structure. Estrogen-dependent maintenance of white matter integrity delays downstream degeneration and enables prefrontal hyperconnectivity that supports cognitive resilience during aging. These results shed new light on sex differences in brain aging and highlight estrogen-sensitive pathways as potential targets for promoting healthy cognitive aging.

## Acknowledgements

We thank the *In Vivo* Imaging facility of the Instituto de Neurociencias for data acquisition. We also thank the Animal Experimentation Service of the Universidad Miguel Hernández for the care and housing of the animals, and for providing a space to carry out the behavioral testing. We thank Dr. S. Jurado and Dr. B. Strange for fruitful discussions. SDS was supported by the Spanish Ministerio de Ciencia e Innovación, Agencia Estatal de Investigación (PID2021-128909NA-I00 and CNS2023-14488), by the Programs for Centres of Excellence in R&D Severo Ochoa (CEX2021-001165-S), by the Generalitat Valenciana through a Subvencion para la contratación de investigadoras e investigadores doctores de excelencia 2021 (CIDEGENT/2021/015 and CIESGT/2025/06), and by the Fundacion Pasqual Maragall (ref. 2023-1296). SC was supported by the Spanish Ministerio de Ciencia e Innovación and the Agencia Estatal de Investigación (PCI2024-153491 funded by MICIU/AEI/10.13039/501100011033) and the European Union through the ERA-NET NEURON framework project IBRAA, as well as by projects PID2021-128158NB-C21 and PID2024-162400OB-C21 (MICIU/AEI/10.13039/501100011033), and the Generalitat Valenciana and Next Generation EU through the Prometeo Excellence Grant (CIPROM/2022/15). ME was supported by a Junior Leader Fellowship of the “la Caixa” Foundation with Project-ID LCF/BQ/PI23/11970039.

## Methods

### Animals and experimental design

Animals were housed 2–4 per cage (Type-IV; Ehret, Germany) under a 12-hour light/dark cycle, with *ad libitum* access to food and water. Animal health and body weight were regularly monitored at the end of each experimental session to ensure welfare. All animals were purchased from Janvier Labs (Le Genest-Saint-Isle, France), and were maintained under the same housing conditions in the animal facility. All experimental procedures were approved by the Animal Care and Use Committee of the Instituto de Neurociencias de Alicante (Spain) and were conducted in compliance with Spanish (Law 32/2007) and European regulations (EU Directive 86/609, EU Decree 2001-486, and EU Recommendation 2007/526/EC).

Twenty-nine Wistar rats (14f/15m) were used in a longitudinal study spanning two years, from 35 to 731 postnatal days (PND). Diffusion-weighted MRI (dw-MRI) data were acquired at thirteen time points throughout the lifespan: 35, 63, 91, 119, 161, 210, 280, 364, 420, 483, 546, 609 and 731 PND, for a total of 215 experiments. Resting-state functional MRI (rs-fMRI) acquisitions were performed at 91, 210, 364, 546 and 731 PND, for a total of 67 experiments. Experimental design with number of animals for each timepoint is illustrated in Fig.1a. At the final time point (731 PND), the rats that survived until the end of the study (5f/3m) underwent the Morris Water Maze test to assess spatial learning and memory performance, then were perfused for histological analysis. A separate subset of 6 females and 6 males aged 35, 90 and 120 PND (two per age) was also used for histological analysis. An extra cohort of Wistar rats aged 730 PND (3f/3m) from a separate batch was included for additional Morris water maze analyses, since not all animals from the longitudinal experiment could complete the test. This extra cohort also underwent rotarod testing at 1.5 (5f/5m) and 2 (3f/3m) years of age. Experimental design is illustrated in Fig.3g for behavior, and in Fig.4c for histology. To further characterize the MRI findings, electrophysiological experiments were performed in a separate cohort of Wistar rats. A total of 18 experiments were conducted across 6-month-old (4 females, 4 males) and 1-year-old animals (4 females, 6 males) (±1 month). A small number of recordings were excluded due to technical issues or atypical responses, reflecting minor deviations in electrode placement. Experimental design is illustrated in Fig. 4f and S6.

To investigate the influence of sex hormones, a second longitudinal experiment was performed with 5 ovariectomized (OVX) female Wistar rats, as well as 5 sham-operated females and 5 males. MRI acquisitions were performed at 120, 180, 400 and 500 PND (±20 days, due to technical constraints related to MRI scheduling), including dw-MRI and rs-fMRI. At 400 and 520 PND, animals (5 OVX, 5 sham males, 2 sham females) underwent Morris water maze test to assess spatial learning and memory performance.

### MRI acquisition

MRI experiments on rats were performed on a 7T scanner (Bruker, BioSpect 70/30, Ettlingen, Germany) using a receive-only phase array coil with integrated combiner and preamplifier in combination with an actively detuned transmit-only resonator. During all acquisitions, animals were anesthetized with isoflurane/O₂ at 3-5% (vol/vol), maintained at 2% vol/vol during anatomical and diffusion sequences and reduced to 1.5% vol/vol during rs-fMRI. Physiological parameters including temperature (37–37.5°C), heart rate, and SpO₂ were continuously monitored, and body temperature was controlled using a water-heating pad. Dw-MRI images were acquired using a spin-echo echo-planar imaging sequence with 16 coronal slices and 20 uniformly distributed diffusion gradient directions, b= 1000 and 2500 s/mm², together with three non-diffusion-weighted images (*b* = 0). Additional parameters were: diffusion time = 15 ms, repetition time (TR) = 5000 ms, echo time (TE) = 26.3 ms, and three repetitions. T2-weighted anatomical images were obtained using a rapid acquisition with relaxation enhancement (RARE) sequence with the following parameters: effective TE = 45 ms, TR = 3000 ms, RARE factor = 8, and four averages. T1-weighted images were obtained with a RARE-VTR sequence with the following parameters: TE = 12 ms, TR = 300 ms, 2 repetitions. The geometry was matched across diffusion and RARE sequences to be the following: FOV = 25 × 25 mm, 16 slices, slice thickness = 1 mm, matrix 110 × 110. Resting-state functional MRI was acquired using a gradient-echo echo-planar imaging sequence with 56 coronal slices and the following parameters: FOV = 25 × 25 mm, slice thickness = 0.5 mm, matrix = 50 × 50, one segment, flip angle = 60°, TE = 15 ms, TR = 2 s and a total of 600 volumes. High-resolution T2-weighted anatomical images optimized for functional analysis were obtained using a RARE sequence with parameters: FOV = 25 × 25 mm, 56 slices, slice thickness = 0.5 mm, matrix = 200 × 200, effective TE = 11 ms, TR = 2 s and RARE factor = 8. Total scan time, including animal positioning, was around 2 h 30 minutes. MRI acquisitions for the OVX cohort were performed using the same scanner and general acquisition protocol described above, except for the following differences: no T1-weighted contrast was acquired, and 450 volumes per acquisition for resting-state functional MRI. In addition, minor modifications to the diffusion protocol were introduced at 400 PND due to technical constraints following an MRI scanner failure. Specifically, the diffusion gradient duration was increased from 3 to 3.2 ms, and the echo time was increased to 26.5 ms. Because of these minor protocol differences, data acquired with different protocols were not directly compared.

### MRI Processing

Dw-MRI raw data was processed using in-house Matlab scripts relying on several standard toolkits. The ANTs (Advanced Normalization Tools) registration framework^67^ was used for all the image registrations. FSL BET (Brain Extraction Tool)^68^ was used to remove the skull from the T1-weighted and dw-MRI data, separately for each contrast. Data were non-linearly registered to the T1-weighted scan to correct for echo-planar imaging distortions using 2D deformation in the x-y plane, then corrected for motion and eddy current distortions using 3D affine registration. For the OXV cohort where T1-weighted scan was not available, T2-weighted was used with comparable results in terms of artefact correction. The shell with b-value of 1000 s/mm² was used to extract diffusion tensor imaging maps, specifically mean diffusivity (MD) and fractional anisotropy (FA). All volumes were used to extract maps of microstructural integrity according to the CHARMED framework^69^, namely, the restricted water fraction (FR), and the angular dispersion of the restricted compartment’s orientation (FD). For each subject, the transformation to the Waxholm rat brain atlas^70^ was calculated with ANTs using both FA and T2-weighted contrasts (one optimal for white matter, the other for grey matter). Then, the inverse transform was applied to the atlas parcellation, to bring parcellated ROIs in the space of each individual subject. The average for each microstructural parameter (FA, MD, FR and FD) was then obtained for each region.

For functional MRI analysis, high-resolution T2-weighted MRI images were first corrected for intensity inhomogeneities using bias field correction, then used to construct a study-specific anatomical template serving as a common reference space for all subjects through the ANTs function *buildtemplate*. Resting-state fMRI data were preprocessed by skull stripping, motion correction, spatial smoothing (full width at half maximum 7 mm), intensity normalization, and high-pass temporal filtering. Noise components were identified and removed using independent component analysis (ICA) implemented in FSL MELODIC^71^. Group-level ICA was subsequently performed on all fMRI data from all subjects concatenated over time to identify resting-state networks (RSNs) in the common anatomical template space. The group ICA was performed on all fMRI data from all subjects from the first longitudinal experiment, concatenated over time (multi-session temporal concatenation, FSL). A total of 25 components were extracted, of which 11 were identified as nodes and the rest as noise components. The ICA identified as networks are shown in Fig. S3. All the analyses were performed using custom commands developed in Python, combining ANTs^67^, FSL^68^ and AFNI^72^ software. For the second longitudinal experiment including OVX and sham animals, we applied the same processing but focused on the default mode network only. Finally, brain volume measured as the size of the skull-stripped T2W image was used to test effect of sex and age using a linear mixed model, including weight as covariate.

### Microstructural MRI statistical analysis

To characterize lifespan trajectories of white and gray matter microstructural parameters (FA, MD, FD, and FR), we first used the Bayesian Information Criterion to identify the model that best balanced goodness of fit and parsimony. The typical trajectory of microstructural parameters is with age characterized by an initial phase of maturation (increasing anisotropy, decreasing diffusivity), followed by a phase of maturity and aging (decreasing anisotropy, increasing diffusivity), as described in previous aging studies^37,73^. Segmented regression is a flexible alternative to traditional quadratic approaches, and unlike them, this model does not assume a symmetrical change around the peak, allowing us to independently estimate the rate of maturation and the rate of decline. We compared then segmented regression with a more parsimonious linear model. In regions of interest (ROIs) where a segmented model was supported, representing the majority, we focused on two key features: (i) the age at which the trajectory changed direction, hereafter referred to as the breakpoint or inflection point; and (ii) the slope, representing the rate of change after this point. Accordingly, for each ROI, lifespan trajectories were modeled using segmented regression to estimate both the breakpoint and the post-breakpoint slope. To test the effect of sex and in line with previous work^16^, we considered two segmented regression approaches: i) a model including both sexes with a single breakpoint shared between sexes; and ii) a sex-specific model fitted independently to males and females, allowing estimation of sex-specific breakpoints. All statistical analyses for this stage were conducted in R v4.3.2 using the *segmented* package. Outliers were identified using the Robust Regression and Outlier removal method (Q = 1%) in GraphPad Prism 8.0.1 (GraphPad Software, San Diego, CA, USA). Differences between sexes were evaluated using paired t-tests. In addition, to investigate spatial trends (anterior–posterior, A–P) in the post-breakpoint slope across white matter tracts, we tested the relationship between the slope estimated from the most explanatory model for each ROI and the A–P coordinates of the ROI-wise geometric centers (i.e., the average position of each ROI within the brain) using Pearson correlation. Human diffusion MRI data from the cohort described in ^16^ were reanalyzed to investigate age-related trajectories of grey matter MD. Diffusion images were preprocessed as in the original study, and the b0 image was registered to the Desikan–Killiany atlas^74^. Atlas labels were then transformed back into each subject’s native diffusion space to extract regional MD values from grey matter regions. For each ROI, MD was modeled as a quadratic function of age separately in males and females, following the same framework used for white matter parameters in the original study. In regions showing a significant quadratic relationship with age, the trajectory breakpoint, interpreted as the age at which the parameter reverses its trend, was calculated from the fitted model. Sex differences in breakpoint age were then quantified by comparing the estimated values between males and females.

### Functional MRI statistical analysis

Dual regression^75^ was performed on the group-level component maps for the first longitudinal experiment. In the first step, subject-specific time courses were extracted for each node by regressing the group spatial maps onto each subject’s fMRI data. These time courses were then variance-normalized and, in the second step, regressed back onto the individual fMRI data to obtain subject-specific spatial maps for each node. The resulting spatial maps are sensitive to both signal amplitude and spatial configuration. To investigate functional interactions between the identified RSNs, a network-level analysis was performed using FSLNets (v0.6). Subject-specific time courses extracted during the first stage of the dual regression served as input. Group-ICA components were used as network nodes, and the corresponding time courses were further denoised and high-pass filtered to ensure signal stability. For each subject and time point, connectivity matrices were generated by estimating pairwise interactions between nodes using both full correlation, reflecting total connectivity, and L2-regularized partial correlation (ρ = 0.01). Partial correlation was prioritized to estimate direct interactions between networks while accounting for indirect influences from other nodes. Fisher Z-transformed correlation coefficients were then analyzed using a linear mixed-effects model framework to assess age-related changes in edge strength.

In addition, group-level ICA maps were used as spatial masks to extract mean parameter estimates (raw beta values) from each subject’s stage-2 spatial maps for each RSN. These values were used to characterize lifespan trajectories of functional connectivity using linear mixed-effects models for each RSN. Fixed effects included age, sex, their interaction, and a quadratic age term to capture non-linear trends. Multiple comparison across voxels were accounted for by using a permutation-based framework. Subject identity was included as a random effect to account for repeated measures and inter-individual variability. The quadratic model was selected over a linear alternative based on the Bayesian Information Criterion, reflecting improved model fit and parsimony. Statistical analyses were performed in R (v4.3.3) using the *lme4* and *lmerTest* packages, and using the *randomise* function in FSL.

For the second longitudinal experiment including OVX and sham animals, we performed a seed-based analysis focused on the default mode network, defined as the region-of-interest mask of the first, denser longitudinal analysis. For each subject and time point, the mean fMRI time series was extracted from the DMN mask. Dual regression was applied as before. Beta values were modeled as a function of age, group, and their interaction, with animal identity included as a random intercept in *randomise*, while correcting for multiple comparison across voxels.

### Combined functional and microstructural analysis across timepoints

To investigate the temporal relationship between microstructural and functional changes, we performed a cross-modal analysis across time points. For all pairs of time points, the mean FA within the white matter skeleton was correlated with subject-specific beta values. This resulted in a matrix of correlation analyses spanning all combinations of structural and functional time points. For each pair, the number of significant correlations was quantified, generating a summary matrix reflecting the temporal distribution of cross-modal associations. To assess the directionality of these relationships, namely, whether early microstructural measures predict later functional connectivity, the reverse, or whether both are associated at the same time point, the symmetry of the resulting matrix was evaluated using a Wilcoxon signed-rank test comparing values above and below the diagonal.

### Monte Carlo Simulation of 2D Microstructural substrate

A two-dimensional gray matter substrate was generated using an in-house Python (v3.11) framework designed to model cellular microstructure. The package creates cell geometries as polygonal approximations of cell bodies and processes using the *Shapely* library, allowing flexible generation of diverse cell populations through pre-defined configuration files. For this study, populations of neurons, microglia, and astrocytes were generated, including both healthy and pathological variants^76^. Neurons were modeled with a soma, a primary axon extending from the cell body, and a set of dendrites radiating outward; pathological neurons were defined by a reduced dendritic arbor. Microglia were represented by smaller somas with distinct processes extending radially, whereas pathological microglia were larger and lacked visible processes. Astrocytes were modeled with comparatively larger somas occupying much of the cell extent, with processes extending outward; pathological astrocytes were defined primarily by increased size. The relative proportion of these populations was 5:3:1 for neurons, astrocytes and microglia, respectively. These three cell populations were distributed within a square domain approximately 2 mm × 2 mm using a Poisson disk sampling approach to generate candidate positions, followed by a packing step to achieve a dense and realistic representation of brain gray matter. The same spatial distributions were reused for the corresponding pathological variants to ensure consistency. Diffusion within the substrate was simulated by initializing 100,000 particles distributed across both intra- and extracellular compartments. Particles underwent random walks with fixed step sizes, reflecting off cellular boundaries when encountered, and were constrained to remain within their initial compartment to prevent exchange between intracellular and extracellular spaces. Step size and number of steps were selected to approximate physiological water diffusion over experimentally relevant timescales.

To generate the simulated diffusion-weighted MRI signal, the acquisition protocol was replicated within the simulation, including matching b-vector orientations, b-values, and diffusion times. The diffusion-weighted signal was then computed by discretely integrating the trajectories and magnetic phase accumulation of the ensemble of particles, yielding a simulated dw-MRI signal consistent with the experimental design. To assess changes between baseline and pathology, the percentage change was calculated for each subject and tested against zero using a one-sample Wilcoxon signed-rank test. Resulting p-values were corrected for multiple comparisons using the Benjamini–Hochberg procedure.

### Behavioral testing

Behavioral tests were conducted to evaluate motor coordination and spatial memory. Motor coordination and balance were assessed using an accelerating rotarod apparatus (80-mm diameter drum; Harvard Apparatus, model LE8355). Rats were placed on the rotating rod and allowed to walk freely. The training phase consisted of two consecutive days, during which the rod rotated at a constant speed of 4 rpm. Animals performed four training sessions per day, each lasting up to 5 min, with 10-min inter-session intervals. Animals that fell were immediately returned to the rod and allowed a few seconds to re-acclimate before resuming the trial. The testing phase was conducted over the following three consecutive days using an accelerating rotarod protocol. Each day included three trials separated by 10-min rest intervals. During each trial, rod speed increased from 4 to 40 rpm over 5 min. Latency to fall was automatically recorded using pressure sensors and used as an index of motor coordination and motor learning. Performance across testing sessions and days was analyzed as a measure of motor coordination and motor learning. Latency to fall during the testing was analyzed using a linear mixed-effects model with day, sex, and their interaction as fixed effects, animal identity as a random effect, and body weight as a covariate. All statistical analyses were conducted in R v4.3.2 using the *lme4* and *lmerTest* packages. Data are reported as mean ± SEM.

Spatial learning and memory were assessed using the Morris Water Maze to evaluate sex-related differences. Testing was conducted over five consecutive days, with four trials per day. The escape platform was randomly relocated at the start of each day. The first trial of each day assessed long-term memory, requiring animals to recall the platform location from the previous day, while the subsequent three trials evaluated short-term learning of the new platform location. Performance was quantified as the latency (seconds) to locate the hidden platform in each trial. Testing was conducted in a circular white pool (180 cm diameter, 60 cm depth), filled to capacity with a 10-cm gap from the pool edge. Water was made opaque using non-toxic purple water-based ink and maintained at 22 ± 0.5 °C. The escape platform (10 cm diameter) was submerged 2 cm below the water surface at a different location each day. Four distal extra-maze cues were positioned around the room and remained constant throughout testing.

Behavioral data were analyzed using AnimalTA^69^. To quantify learning, the change in latency across days was calculated separately for long-term and short-term memory. For long-term memory, the first trial of each day was used to compute the learning slope across days. For short-term memory, the mean latency of the subsequent three trials (trials 2–4) was calculated for each day, and the slope of these means across days was used as a measure of short-term learning. These slopes were then compared between sexes for each task using unpaired t-test. To ensure sufficient sample size at 2 years of age, the analysis included animals from the longitudinal cohort that survived to this age (4 females, 2 males), combined with additional age-matched animals from the separate cohort (3 females, 3 males). Multiple linear regression confirmed no significant effect of cohort (p=0.60), validating the pooling of animals for analysis.

### Electrophysiological recordings

Rats were intraperitoneally anesthetized with urethane (1.4 g/kg), with supplemental doses (one-fifth of the initial dose) administered as needed to maintain the absence of reflexes. Animals were then secured in a stereotaxic frame, and physiological parameters were monitored throughout: animals breathed a mixture of air and oxygen (0.8 L/min), and body temperature was maintained at 37 °C using a water-heating pad. At the end of the experiments, animals were sacrificed, and brains were processed to verify electrode placement. Surgical and stereotaxic procedures were performed as previously described^77^. A concentric bipolar stimulating electrode (WPI, London UK) was positioned in the fimbria (from bregma AP −1.5, ML 0.4, DV 3,2 mm), according to rat brain atlas^78^. Two 32-channel silicon probes (single shank, 100 μm spacing; Neuronexus Technologies) were lowered into the intermediate hippocampus and medial prefrontal cortex (covering prelimbic and infralimbic areas) (coordinates from bregma: hippocampus AP −4.4, ML 2.6, DV 3.5 mm; and prefrontal cortex AP 3.4, ML 0.5, DV 5 mm). Probes were pre-coated with DiI for postmortem confirmation of placement. A silver chloride wire in contact with the neck skin served as ground. Electrophysiological signals were high-pass filtered at 0.1 Hz, amplified, and digitized at 10 kHz using Multi Channel Systems hardware and software. Evoked potentials were recorded in response to electrical stimulation delivered via a pulse generator and current source (STG2004, Multichannel Systems, Reutlingen, Germany). A stimulus-response input-output (I/O) protocol consisted of single biphasic pulses (100 µs duration) at increasing intensities (50–1000 µA), delivered every 15 s. The sequence of intensities was repeated four times. Detailed protocol here: 10.17504/protocols.io.4r3l2dprjg1y/v1.

Neuronal firing in response to the stimulation protocol was quantified as the amplitude of population spikes in CA1 and the amplitude of field excitatory postsynaptic potentials in prefrontal cortex. Statistical analyses of evoked responses were performed using GraphPad Prism 8.0.1 (GraphPad Software, San Diego, CA, USA). For each age group (6 months and 1 year), a two-way repeated-measures ANOVA was used to assess the effects of stimulation intensity, sex, and their interaction on amplitude in CA1 and prefrontal cortex. When significant interactions were detected, Sidak’s multiple comparisons test was applied to assess differences between groups.

In addition, correlation analyses between CA1 and prefrontal cortex were performed. Activity propagation from HC to PFC was quantified as the amplitude of the evoked potential in the PFC divided by the simultaneously recorded PS in CA1, referred to as the propagation ratio. Separate unpaired t-tests were used to assess the effect of age on propagation ratio in males and females.

### Immunofluorescence staining of rat tissue

Rats undergoing immunofluorescence during maturation were intracardially perfused with 0.9% saline + Heparin 0.1% (v/v) and 4% paraformaldehyde (PFA; w/v). For the cohort of young animals (30–120 PND), the left hemisphere of brains was embedded in 3% agarose/PBS (Sigma-Aldrich, Madrid, Spain) and cut on a vibratome (VT 1000S, Leica, Wetzlar, Germany) into 50 μm-thick serial sagittal sections. Sections underwent citrate antigen retrieval, permeabilization, and blocking for 2 h at room temperature in 4% bovine serum albumin and 2% goat serum (Sigma-Aldrich). A total of five antigen labelings were performed: myelin, neurofilaments, neuronal nuclei, microglia and astrocytes. Three sagittal slices, that englobed the 15 mentioned ROIs, were selected per rat for each labelling: a lateral, a medial and a central one. The slices were then incubated for one night at 4°C with primary antibodies for Iba-1 (1:1000, Wako Chemicals, Osaka, Japan), GFAP (1:1000, Sigma Aldrich, Madrid, Spain), neurofilament (1:200, Abcam, Cambridge, United Kingdom), MBP (1:250, Merck Millipore, Darmstadt, Germany) and NeuN (1:250, Merck Millipore, Darmstadt, Germany), to label microglia, astrocytes neuronal processes, myelin and neuronal nuclei, respectively. Sections were then incubated with secondary antibody (Alexa Fluor™ 594, 1:500) for 2 h at room temperature in a humid chamber. Following counterstaining with DAPI, slides were coverslipped with Mowiol. Sections were imaged using a Leica DM400 fluorescence microscope equipped with a QICAM Qimaging camera 22577 (Biocompare, San Francisco, USA) and Neurolucida software. Images were acquired with a 2.5× objective, covering 15 ROIs per section: 6 white matter (splenium and genu of corpus callosum, fimbria, anterior commissure, cingulum, and cerebellar peduncle) and 7 belonging to grey matter (prelimbic cortex, association cortex, hypothalamus, hippocampus, amygdala, nucleus accumbens and substantia nigra). The staining intensities was quantified using ICY Software within an area of 100 μm². Repeated-measure ANOVA (factors: age and ROI) performed on the mean intensity of immunohistological staining for neurons (NeuN and neurofilaments) and myelin (MBP), and quantitative cell count of glia population (Iba-1 for microglia and GFAP for astrocytes) on 7 WM ROI revealed that only MBP intensity presented a statistically significant variation with time (p=0.013).

The brains from rats from the longitudinal experiment at the final time point (PND 731) were post-fixed overnight and cryoprotected in 30% sucrose before embedding in OCT (Tissue-Tech) and storage at −80 °C. Cryosections (30 μm) received citrate antigen retrieval and permeabilized and blocked for 2 h with 5% normal goat serum (Merck) with 0.5% Triton-X-100 (Merck), in PBS. Primary antibodies were applied overnight at 4 °C in a humid chamber and included the following: GFAP (Abcam, 1:1000; Ab4674); Iba1 (Wako, 1:5000; 019-19741); NeuN (Merck, 1:1000;MAB377), GAD67 (Merck, 1:500; MAB5406). Secondary antibodies were applied for 2 h at room temperature in a humid chamber (1:500, Alexa Fluor™ 488 A-11039,CF™ 555 antibody SAB4600069, Alexa Fluor™ 647 A21236). Following counterstaining with DAPI, slides were coverslipped with VECTASHIELD® HardSet™ Ref H-1400. Detailed protocol here: 10.17504/protocols.io.kxygx83j4v8j/v1 for NeuN/Iba1/GFAP and here: 10.17504/protocols.io.81wgbox8qlpk/v1 for GAD67. Sections were imaged on a Leica Thunder Imager microscope (Leica DMI8, Objetive 20x, Dimesion Size 1024×1024 pixels). Selected regions included frontal areas (prefrontal cortex, motor cortex), parietal association cortex, retrosplenial cortex, CA1. Cell counts were calculated using Imaris 10.2. The proportion of inhibitory neurons was calculated by co-localizing GAD67 with NeuN. Unpaired t-tests were used to compare across sexes after averaging multiple repetitions for each region of interest.

### Statistical considerations

Multiple comparison correction was applied to the longitudinal and all voxel-wise analyses, which were sufficiently powered, as detailed in each section. For subsequent experiments with smaller sample sizes, no correction was applied unless explicitly mentioned, as these analyses were hypothesis-driven and limited to a much smaller number of comparisons.

