## Supplementary material for "Longitudinal MRI reveals hormone-dependent brain remodeling supporting preserved cognition in aged females"

**Extended data**

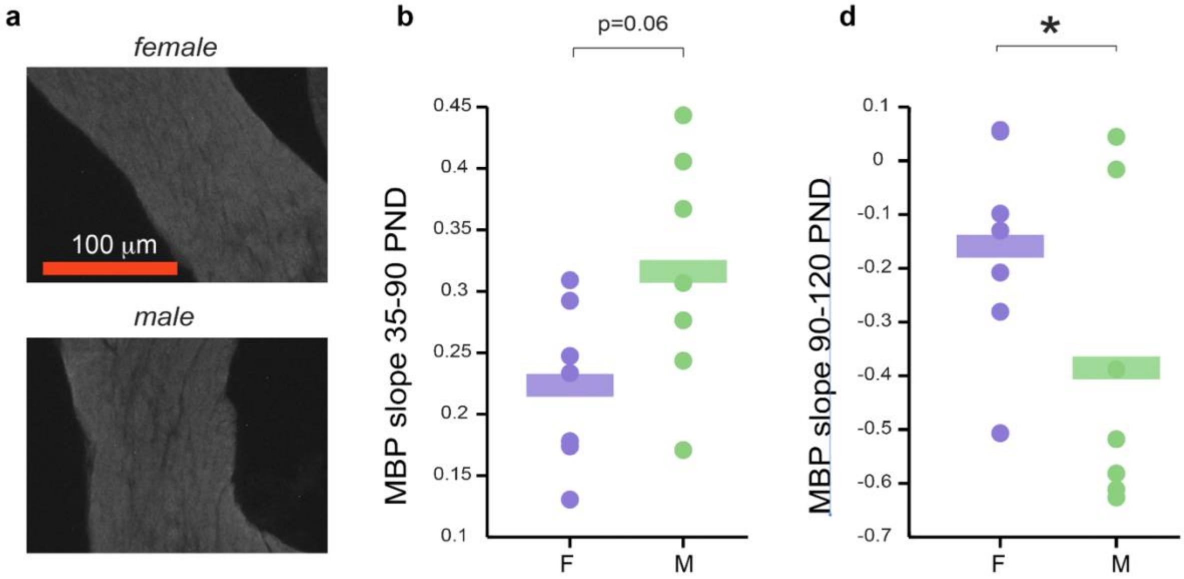


***Figure S1: Myelin basic protein quantification in white matter. a.*** *Representative immunostaining images of myelin basic protein (MBP) in white matter tracts in females (upper image) and males (bottom images). Red scale bar: 100 μm.* ***b.*** *MBP intensity over postnatal development in females (purple) and males (green), showing the trajectory up to 120 PND.* ***c.*** *Slope of MBP maturation for all white matter ROIs. Males (green) exhibited a trend toward faster initial maturation compared to females (purple) (paired t-test, p = 0.06).* ***d.*** *Slope of MBP decline, showing a significantly steeper decrease in males (green) compared to females (purple) (paired t-test, p = 0.032). Abbreviations: F = female; M = male.*

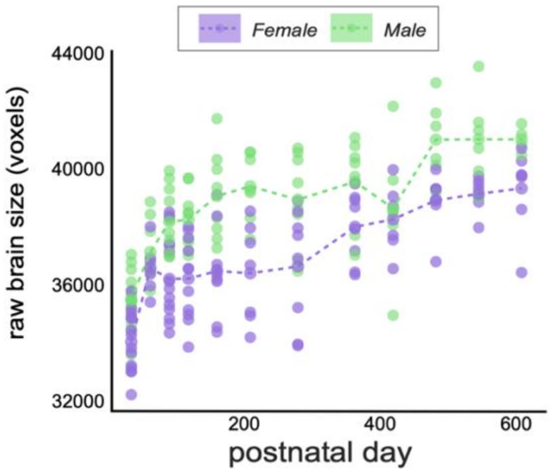

***Figure S2*: Brain size across the lifespan.** *Brain size expressed in voxels for all animals at different post-natal days. In the linear mixed model there was no main effect of sex, but a significant interaction between sex and age (p < 0.001), as well as a significant main effect of age (p = 0.003). Females are shown in purple and males in green.*

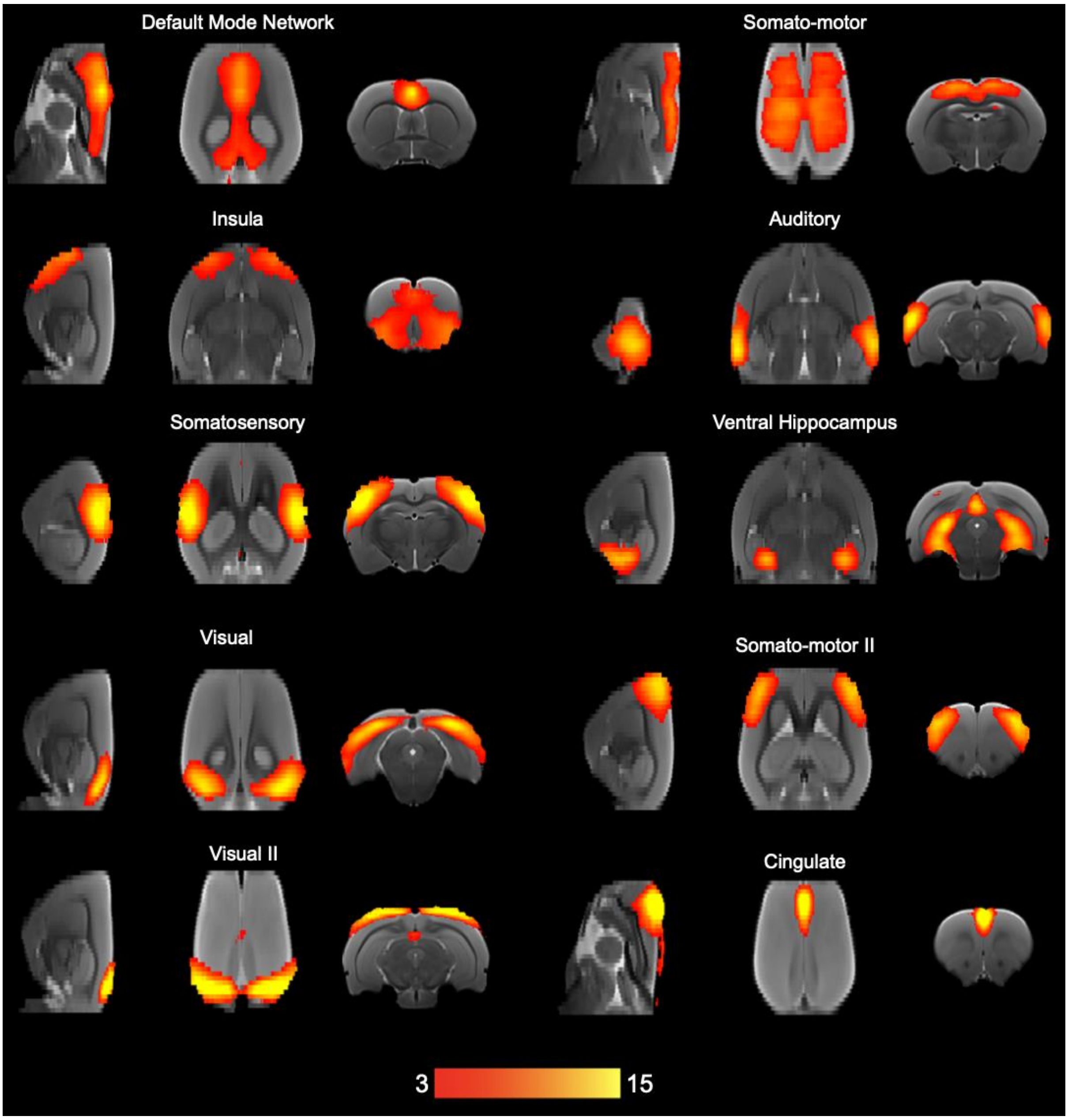


***Figure S3****: ICA components using the same nomenclature used in the consensus paper by Grandjean et al. 2020.*

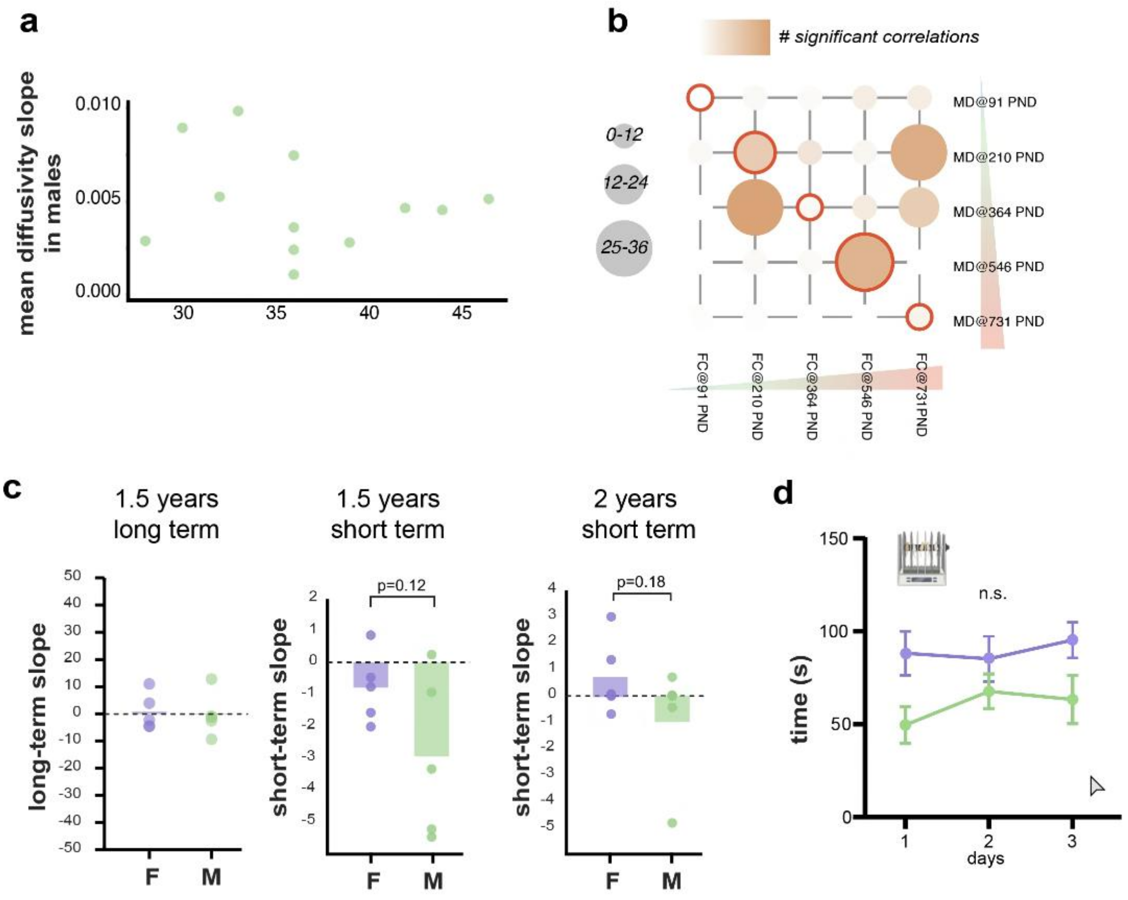


**Figure S4: a.** *White matter MD slope in males plotted as a function of the anteroposterior coordinate. The association is not significant, as assessed by a Pearson correlation test* ***b.*** *Cross-modal correlations between MD and fMRI alterations in grey matter (Wilcoxon sign rank test, p=0.14).* ***c.*** *Long-term slope of water maze task at 1.5 years (right), short-term slope at 1.5 years (center) and long-term slope at 2 years of age (right), plotted separately for males and females.* ***d.*** *Time on the rotarod across days, with no sex differences after adjusting for animal weight (females in purple, males in green).*

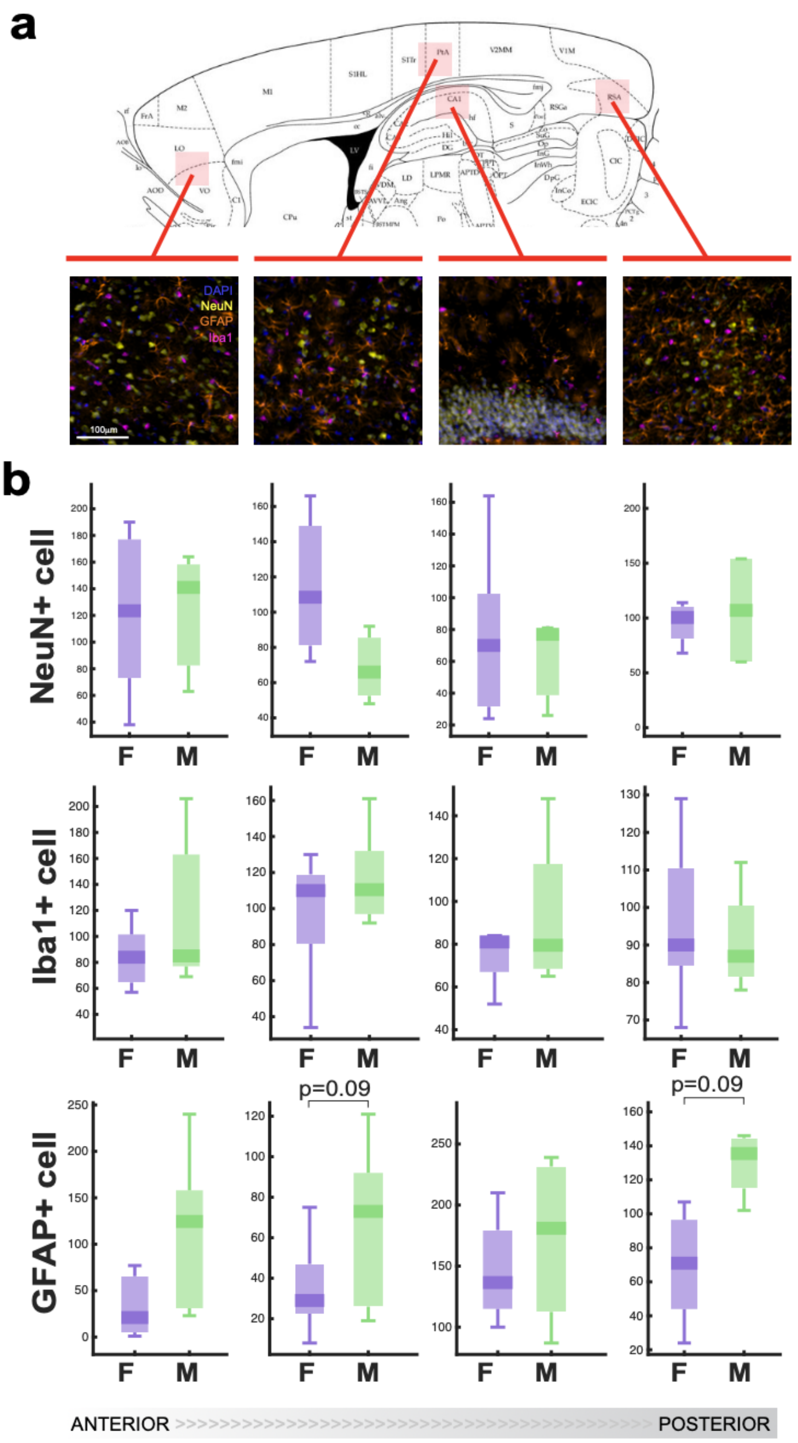


***Figure S5: a.*** *Positioning of ROIs for histology.* ***b.*** *NeuN+, Iba1+ and GFAP+ cell count for different ROIs in the lateral histological slice, plotted separately for males and females. No significant differences are observed for NeuN or Iba1, while a trend of increased GFAP count in males (p=0.09) is reported in the retrosplenial and parietal association cortices.*

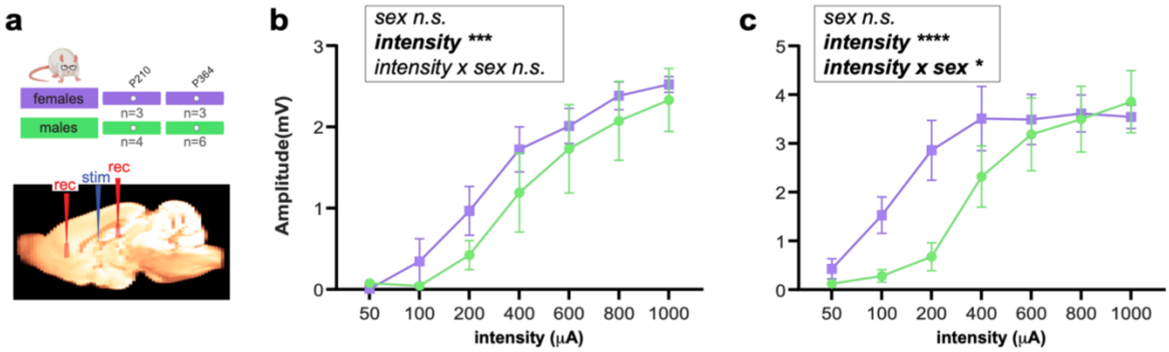


***Figure S6: Evoked response in hippocampus at different ages. a.*** *At 6 months of age, amplitude responses (mV) significantly increase with progressive stimulation intensity (μA) (p = 0.0004), with no effect of sex or sex × intensity interaction.* ***b.*** *At 1 year of age, amplitude responses (mV) also increase with progressive stimulation intensity (μA) (p < 0.0001); and a significant sex × intensity interaction was observed (p = 0.049), while no main effect of sex was detected. Females are shown in purple, males in green. Data are mean ± SEM.*

**Supplementary Table 1.** Preferred model describing MRI parameters trend with age for white matter regions.

| White matter regions | FA | RF | MD |
|---|---|---|---|
| Olfactory tract | Segmented regression | Segmented regression | Segmented regression |
| Nigrostriatal bundle | Segmented regression | Segmented regression | Segmented regression |
| Medial forebrain bundle | Segmented regression | Segmented regression | Segmented regression |
| Internal capsule | Segmented regression | Segmented regression | Linear |

| | | | |
|---|---|---|---|
| (cerebral peduncle) | | | |
| Corpus callosum 1 | Segmented regression | Segmented regression | Linear |
| Corpus callosum 2 | Segmented regression | Segmented regression | Segmented regression |
| Optic tract | Segmented regression | Segmented regression | Linear |
| Cingulum | Segmented regression | Segmented regression | Segmented regression |
| Fornix | Segmented regression | Segmented regression | Linear |
| Fimbria | Segmented regression | Segmented regression | Segmented regression |
| Fasciculus retroflexus | Linear | Segmented regression | Segmented regression |
| Anterior comissure | Segmented regression | Segmented regression | Segmented regression |

**Supplementary Table 2.** Preferred model describing MRI parameters trend with age for grey matter regions.

| **Grey matter regions** | **MD** | **RF** | **FD** |
|---|---|---|---|
| Brainstem | Linear | Linear | Segmented regression |
| Rest of Midbrain 1 | Linear | Linear | Segmented regression |
| Rest of Midbrain 2 | Linear | Segmented regression | Linear |

| | | | |
|---|---|---|---|
| Oculomotor Cortex | Segmented regression | Segmented regression | Segmented regression |
| Compacta Substantia Nigra | Segmented regression | Segmented regression | Linear |
| Entorhinal Cortex | Segmented regression | Segmented regression | Segmented regression |
| Visual Cortex | Segmented regression | Segmented regression | Linear |
| Colliculus | Segmented regression | Segmented regression | Linear |
| Subiculum | Segmented regression | Segmented regression | Linear |
| Retrosplenial Cortex | Linear | Segmented regression | Segmented regression |
| Central Gray | Linear | Linear | Linear |
| Reticular Formation | Segmented regression | Segmented regression | Segmented regression |
| Rest of Midbrain 3 | Segmented regression | Segmented regression | Linear |
| Molecular and Granular Layer of Dentate Gyrus | Segmented regression | Linear | Linear |
| Reticulata Substantia Nigra | Segmented regression | Segmented regression | Linear |
| Lateral Thalamus | Linear | Segmented regression | Linear |
| Cingulate Cortex | Segmented regression | Segmented regression | Linear |
| Auditory Cortex | Segmented regression | Segmented regression | Segmented regression |
| Piriform Cortex | Segmented regression | Segmented regression | Segmented regression |

| | | | |
|---|---|---|---|
| Ventral Tegmental Area | Linear | Segmented regression | Segmented regression |
| Rest of Forebrain 1 | Segmented regression | Segmented regression | Linear |
| Hypothalamus 1 | Linear | Linear | Segmented regression |
| Rest of Midbrain 4 | Segmented regression | Segmented regression | Linear |
| Orbital Cortex | Segmented regression | Segmented regression | Linear |
| Pre-thalamus | Segmented regression | Linear | Linear |
| Somato-sensory Cortex | Segmented regression | Segmented regression | Segmented regression |
| Thalamus 1 | Segmented regression | Linear | Segmented regression |
| Thalamus 2 | Segmented regression | Linear | Segmented regression |
| Medial Thalamus | Segmented regression | Segmented regression | Linear |
| Amygdala | Segmented regression | Linear | Segmented regression |
| Hypothalamus 2 | Linear | Segmented regression | Segmented regression |
| Parietal Association Cortex | Segmented regression | Linear | Segmented regression |
| Part of Medial Complex of Thalamus | Linear | Linear | Linear |
| Thalamus 3 | Segmented regression | Segmented regression | Linear |

| | | | |
|---|---|---|---|
| Rest of Forebrain 2 | Segmented regression | Segmented regression | Linear |
| Caudate, Putamen, Globus Pallidus 1 | Segmented regression | Segmented regression | Segmented regression |
| Thalamus 4 | Segmented regression | Segmented regression | Linear |
| Thalamus 5 | Segmented regression | Segmented regression | Linear |
| Thalamus 6 | Linear | Segmented regression | Segmented regression |
| Caudate, Putamen, Globus Pallidus 2 | Linear | Segmented regression | Segmented regression |
| Thalamus 7 | Segmented regression | Segmented regression | Linear |
| Insular Cortex | Segmented regression | Segmented regression | Segmented regression |
| Motor Cortex | Segmented regression | Segmented regression | Segmented regression |
| Thalamus 8 | Segmented regression | Segmented regression | Segmented regression |
| Thalamus 9 | Linear | Segmented regression | Segmented regression |
| Hypothalamus 3 | Linear | Segmented regression | Segmented regression |
| Septal Nuclei | Segmented regression | Segmented regression | Linear |
| Rest of Forebrain 3 | Segmented regression | Linear | Linear |
| Rest of Forebrain 4 | Linear | Segmented regression | Segmented regression |

| | | | |
|---|---|---|---|
| Olfactory Tubercle | Linear | Segmented regression | Segmented regression |
| Nucleo Accumbens | Segmented regression | Segmented regression | Segmented regression |
| Infralimbic Cortex | Segmented regression | Segmented regression | Linear |
| Prelimbic Cortex | Segmented regression | Segmented regression | Linear |
| CA1 of Hippocampus | Segmented regression | Linear | Segmented regression |
| Polimorphic Layer of Dentate Gyrus (Hilus) | Segmented regression | Segmented regression | Segmented regression |
